# Crystal Structure and Catalytic Mechanism of Drimenol Synthase, a Bifunctional Terpene Cyclase-Phosphatase

**DOI:** 10.1101/2025.02.11.637696

**Authors:** Kristin R. Osika, Matthew N. Gaynes, David W. Christianson

## Abstract

Drimenol synthase from *Aquimarina spongiae* (AsDMS) is a highly unusual bifunctional sesquiterpene synthase that integrates two distinct, sequential isoprenoid processing activities within a single polypeptide chain. AsDMS catalyzes the class II cyclization of farnesyl diphosphate (FPP) to form drimenyl diphosphate, which then undergoes enzyme-catalyzed hydrolysis to yield drimenol, a bioactive sesquiterpene alcohol with antifungal and anticancer properties. Here, we report the X-ray crystal structures of AsDMS and its complex with a sesquiterpene thiol, which are the first of a terpene cyclase-phosphatase. The AsDMS structure exhibits a two-domain architecture consisting of a terpene cyclase β domain and a haloacid dehalogenase (HAD)-like phosphatase domain, with two distinct active sites located on opposite sides of the protein. Mechanistic studies show that dephosphorylation of the drimenyl diphosphate intermediate proceeds through stepwise hydrolysis, such that two equivalents of inorganic phosphate rather than inorganic pyrophosphate are co-products of the reaction sequence. When the AsDMS reaction is performed in H₂^18^O, ^18^O is not incorporated into drimenol, indicating that the hydroxyl oxygen of drimenol originates from the prenyl oxygen of FPP rather than bulk water. These results correct a mechanistic proposal previously advanced by another group. Surprisingly, AsDMS exhibits unprecedented substrate promiscuity, catalyzing the conversion of substrate mimic farnesyl-*S*-thiolodiphosphate into cyclic and linear sesquiterpene products. Structural and mechanistic insights gained from AsDMS expand the functional diversity of terpene biosynthetic enzymes and provide a foundation for engineering “designer cyclases” capable of generating new terpenoid products.

## Introduction

Terpenoids comprise a diverse and broadly useful class of natural products found in all forms of life, having wide-ranging applications in the pharmaceutical, agricultural, biofuel, and fragrance industries.^1–6^ According to the Dictionary of Natural Products,^7,8^ more than 100,000 terpenoids have been identified to date. Strikingly, the vast number of terpenoid natural products belies their roots in two simple 5-carbon precursors, dimethylallyl diphosphate and isopentenyl diphosphate, which are linked together by prenyltransferases to generate linear isoprenoids such as C_15_ farnesyl diphosphate (FPP) or C_20_ geranylgeranyl diphosphate (GGPP).^9,10^ Linear isoprenoids serve as substrates for terpene cyclases, which catalyze the first committed steps in terpenoid biosynthesis to generate structurally complex carbon skeletons typically containing multiple rings and stereocenters. Terpene cyclases catalyze some of the most complex chemical reactions found in nature, since more than half of the substrate carbon atoms undergo changes in bonding and hybridization during the course of a single enzyme-catalyzed reaction.^11–15^

Canonical terpene cyclases are generally grouped into two classes that exhibit various combinations of α-helical domains designated α, β, or γ.^13,16^ A class I enzyme exhibits α, αβ, or αβγ domain architecture and utilizes a trinuclear metal cluster in the α domain to initiate isoprenoid diphosphate ionization, carbocation generation, and cyclization. A class II enzyme exhibits β, βγ, or αβγ domain architecture and utilizes an aspartic acid in the β domain to initiate carbocation generation and cyclization by protonation of an isoprenoid C=C bond.

Intriguingly, some terpene synthases are bifunctional and catalyze two sequential reactions.^17,18^ For example, abietadiene synthase exhibits αβγ domain architecture and catalyzes the class II cyclization of GGPP at the βγ domain interface to generate copalyl diphosphate, which then undergoes class I cyclization in the α domain to yield abietadiene.^19–21^ In another example, fusicoccadiene synthase exhibits (αα)_8_ architecture and catalyzes a chain-elongation reaction in the prenyltransferase α domain to yield GGPP, which is then channeled to the cyclase α domain for the generation of fusicoccadiene.^22–26^

Even more notable, but quite rare, are bifunctional terpene synthases that combine terpene cyclization with a second, completely distinct activity. A recently discovered example is drimenol synthase from the marine bacterium *Aquimarina spongiae* (AsDMS).^27^ This bifunctional sesquiterpene synthase catalyzes the class II cyclization of FPP to form drimenyl diphosphate, which is then hydrolyzed to yield drimenol (Figure 1). Drimenol is a sesquiterpene alcohol known for its antibacterial, antifungal, antiparasitic, and anticancer activities.^27–29^ Based on amino acid sequence analysis, drimenol synthase consists of a terpene cyclase β domain linked to a haloacid dehalogenase (HAD)-like phosphatase domain; these domains are proposed to assemble such that they share a common active site.^27^

**Figure 1.**
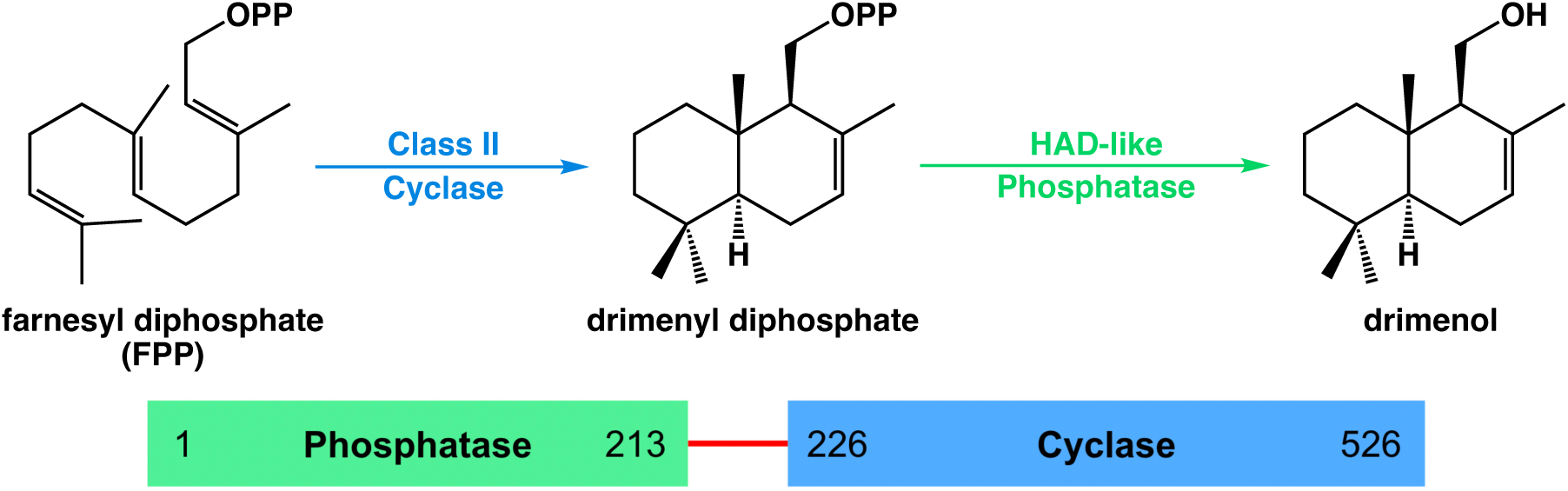
Reaction sequence catalyzed by drimenol synthase from *Aquimarina spongiae* (AsDMS); OPP = diphosphate, HAD = haloacid dehalogenase. The primary structure of the 61-kD AsDMS N-terminal deletion variant prepared for study is shown below; the phosphatase domain is connected to the cyclase domain through a 12-residue linker (red).

Here, we advance our understanding of bifunctional terpene synthases with mechanistic studies and the X-ray crystal structure determination of AsDMS. In contrast with the previously proposed model,^27^ the X-ray crystal structure of AsDMS reveals an alternative assembly of catalytic domains with two distinct active sites. Curiously, an exceptionally unstable primary carbocation intermediate is proposed for the hydrolysis of drimenyl diphosphate to yield drimenol and inorganic pyrophosphate (PP_i_).^27^ Our enzymological studies rule out this pathway and instead support a mechanism in which drimenyl diphosphate undergoes sequential phosphate hydrolysis reactions to yield drimenol and two equivalents of inorganic phosphate (P_i_). Finally, AsDMS cocrystallized with farnesyl-*S*-thiolodiphosphate (FSPP) yields the structure of the complex with farnesyl thiol. Surprisingly, FSPP is sufficiently reactive to enable the AsDMS-catalyzed formation of farnesyl thiol as well as drimenyl thiol over the time course of the cocrystallization experiment, demonstrating AsDMS has a wider substrate scope than previously anticipated.

## Results

### Mechanistic Studies

The conversion of drimenyl diphosphate into drimenol requires hydrolysis of the substrate diphosphate group, but it is not clear whether this occurs through a single reaction yielding inorganic pyrophosphate (PP_i_) or sequential reactions yielding two equivalents of inorganic phosphate (P_i_). The previously proposed AsDMS phosphatase mechanism involves S_N_1 dissociation of the PP_i_ anion to yield a highly unstable primary carbonium ion intermediate, which is quenched by a water molecule from bulk solvent to yield drimenol.^27^ Such a mechanism would be atypical for a HAD-like phosphatase. For example, the HAD-like phosphatase domain of bifunctional human soluble epoxide hydrolase (sEH)^30^ catalyzes the stepwise hydrolysis of isoprenoid diphosphate substrates such as FPP and GGPP to yield the corresponding alcohol and two P_i_ anions.^31^ Since the phosphatase domain of AsDMS is structurally homologous to the phosphatase domain of sEH, we hypothesized that AsDMS would similarly catalyze the stepwise hydrolysis of drimenol diphosphate to yield drimenol and two P_i_ anions.

To test this hypothesis, we utilized the EnzChek pyrophosphate detection assay.^32^ Typically, this assay relies on inorganic pyrophosphatase (IPPase) to generate P_i_ from PP_i_; P_i_ is then utilized as a substrate by purine nucleoside phosphorylase (PNPase) to cleave 2-amino-6- mercapto-7-methylpurine ribonucleoside, which yields the chromophore 2-amino-6-mercapto-7-methylpurine to enable detection and quantification. However, if IPPase were excluded from the assay and if PP_i_ were generated, there would be no measurable signal since PP_i_ cannot be utilized as a substrate by PNPase. Using this assay, we observe a robust signal for P_i_ in the absence of IPPase (Figure S1). This result is consistent only with stepwise hydrolysis reactions yielding two P_i_ anions rather than a single hydrolysis reaction yielding one PP_i_ anion.

The previously proposed phosphatase mechanism of AsDMS suggests that the hydroxyl oxygen atom of drimenol derives from a water molecule.^27^ However, the hydroxyl oxygen atom of drimenol might alternatively derive from the prenyl oxygen atom of FPP. To distinguish between these two possibilities, a sample of AsDMS was lyophilized and resuspended in H_2_^18^O prior to incubation with FPP. Extraction of product drimenol from the reaction mixture and analysis using gas chromatography-mass spectrometry (GC-MS) shows that the molecular weight of drimenol remains unchanged at 222.2 (Figure S2); if ^18^O were incorporated, the molecular weight would increase to 224.2. Therefore, the hydroxyl oxygen of drimenol must derive from the prenyl oxygen of substrate FPP and not a water molecule from bulk solution.

This mechanistic conclusion is also consistent with the generation of drimenyl thiol from FSPP, since the prenyl sulfur atom of FSPP is retained in drimenyl thiol. FSPP might be presumed to be an unreactive substrate analogue due to the substitution of a sulfur atom for the prenyl oxygen atom. However, we considered the potential reactivity of FSPP due to the observation of farnesyl thiol bound in the AsDMS active site in cocrystallization experiments described below. Surprisingly, GC-MS analysis of organic extracts of the reaction mixture after AsDMS is incubated with FSPP overnight reveals the generation of three different hydrocarbon products on the gas chromatogram (Figure S3). Comparisons of chromatographic retention times and mass spectra with those of authentic sesquiterpene alcohol samples suggest that two of the products generated correspond to farnesyl thiol and drimenyl thiol (chromatographic retention times of 10.27 min and 10.72 min, respectively; butterfly plots of mass spectra are recorded in Figure S4A,B). A third peak in the chromatogram with a retention time of 10.64 min could not be conclusively identified by mass spectrum but exhibits a fragmentation pattern that appears to be consistent with a sesquiterpene thiol; this product is generated in equal proportion to drimenyl thiol and likely represents a prematurely quenched monocyclic product (Figure S4C). The retention of the prenyl sulfur atom of FSPP in drimenyl thiol is consistent with the stepwise hydrolysis of the thiodiphosphate group by the phosphatase domain of AsDMS.

Finally, while metal ions are not required for the catalytic mechanism of a class II cyclase (although Mg^2+^ may be required for substrate binding^33^), Mg^2+^ is essential for the dephosphorylation reaction catalyzed by a HAD-like phosphatase.^34,35^ To confirm the expected requirement for Mg^2+^ in catalysis by AsDMS, we measured activity with FPP in the absence of metal ion and in the presence of Mg^2+^, Ca^2+^, Mn^2+^, Co^2+^, Ni^2+^, and Zn^2+^. The generation of drimenol was observed only in the presence of Mg^2+^.

### Crystal Structures

The 2.2-Å resolution crystal structure of AsDMS complexed with the benzyltriethylammonium cation (BTAC), P_i_, and Mg^2+^ reveals that the HAD-like phosphatase domain and the terpene cyclase β domain each contain a distinct active site (Figure 2A). This didomain architecture revises the model previously advanced by another group in which the two catalytic domains share a common active site.^27^ Two essentially identical monomers are found in the asymmetric unit of the *C*2 unit cell, related by a pairwise root-mean-square deviation (rmsd) of 0.615 Å for 457 Cα atoms. Analysis of the crystal structure using PISA^36^ suggests that AsDMS forms a biological dimer. However, mass photometry indicates that AsDMS is a monomer in solution (Figure S5), so the apparent dimerization evident in the crystal structure may be an artifact of crystal packing interactions.

**Figure 2.**
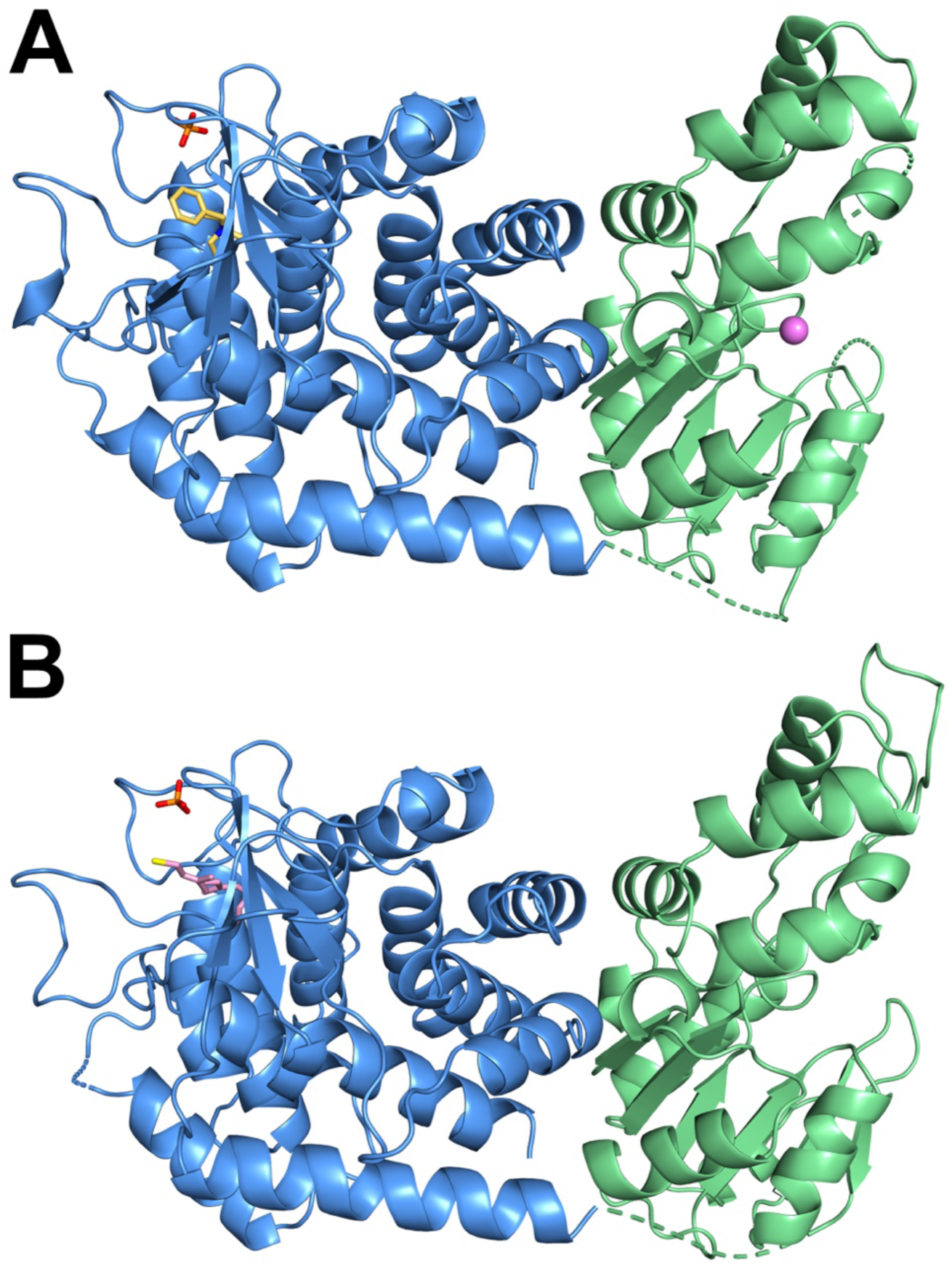
(A) Crystal structure of AsDMS complexed with BTAC, P_i_, and Mg^2+^. The cyclase and phosphatase domains are blue and green, respectively. BTAC (yellow) and P_i_ (P = orange, O = red) are bound in the cyclase active site, while Mg^2+^ (violet sphere) is bound in the phosphatase active site. Notably, the two active sites are on opposite sides of the protein. Dotted lines indicate disordered segments. (B) Crystal structure of AsDMS complexed with farnesyl thiol. Color-coding is identical to that in (A), with farnesyl thiol shown in pink.

The cyclase β domain of AsDMS is formed by a bundle of 12 α-helices and closely resembles the β domains found in other class II terpene cyclases. Intriguingly, however, all other canonical class II terpene cyclases exhibit βγ or αβγ domain architecture, with the active site located at the βγ domain interface.^12,14,37^ In view of the sequence identity and structural homology between the β and γ domains,^38^ which are oriented toward each other in βγ assemblies, the γ domain is thought to be the result of gene duplication and fusion with the β domain early in the evolution of terpene synthases. The domain architecture of AsDMS is quite different in that the phosphatase and cyclase active sites are oriented away from one another (Figure 2). Consequently, the class II cyclase active site in the β domain of AsDMS is more solvent-exposed compared with the cyclase active site in a βγ or αβγ domain assembly, e.g., as observed in squalene-hopene cyclase or copalyl diphosphate synthase, respectively.^39–41^ The protein structure most closely related to the β domain of AsDMS is that of the meroterpene cyclase MstE, which catalyzes the class II cyclization of the ubiquinone-like substrate 5- geranylgeranyl-3,4-dihydroxybenzoate.^42^ Analysis using DALI^43^ indicates that these two proteins are related by 12% sequence identity, a Z score of 23.9, and an RMSD of 3.0 Å for 347 residues (Figure S6).

The β domain active site is located in a hydrophobic pocket situated in the middle of the helix bundle (Figure 3A). A single P_i_ anion binds at the top of the active site, stabilized by ideally-oriented^44^ hydrogen bonds with R368, R370, and R415. These residues may also interact with the diphosphate group of FPP to hold the substrate in a productive conformation for cyclization. BTAC binds more deeply in the hydrophobic pocket, such that its quaternary ammonium cation makes electrostatic interactions with the catalytic DXDT motif at the base of the pocket (the second aspartic acid in this motif is the general acid catalyst that initiates the cyclization cascade). BTAC has been utilized as a partial analogue of a carbocation intermediate in structural studies of class I sesquiterpene cyclases, such as *epi*-isozizaene synthase,^45^ and is a useful indicator of possible enzyme-substrate interactions. The structure of the AsDMS-BTAC complex is the first of a BTAC complex with a class II cyclase, and the binding of BTAC may similarly exemplify possible enzyme-substrate interactions. For example, cation−π interactions observed between the quaternary ammonium group of BTAC and the aromatic side chains of F271, F308, and Y417 may similarly stabilize carbocation intermediates in catalysis by AsDMS.

**Figure 3.**
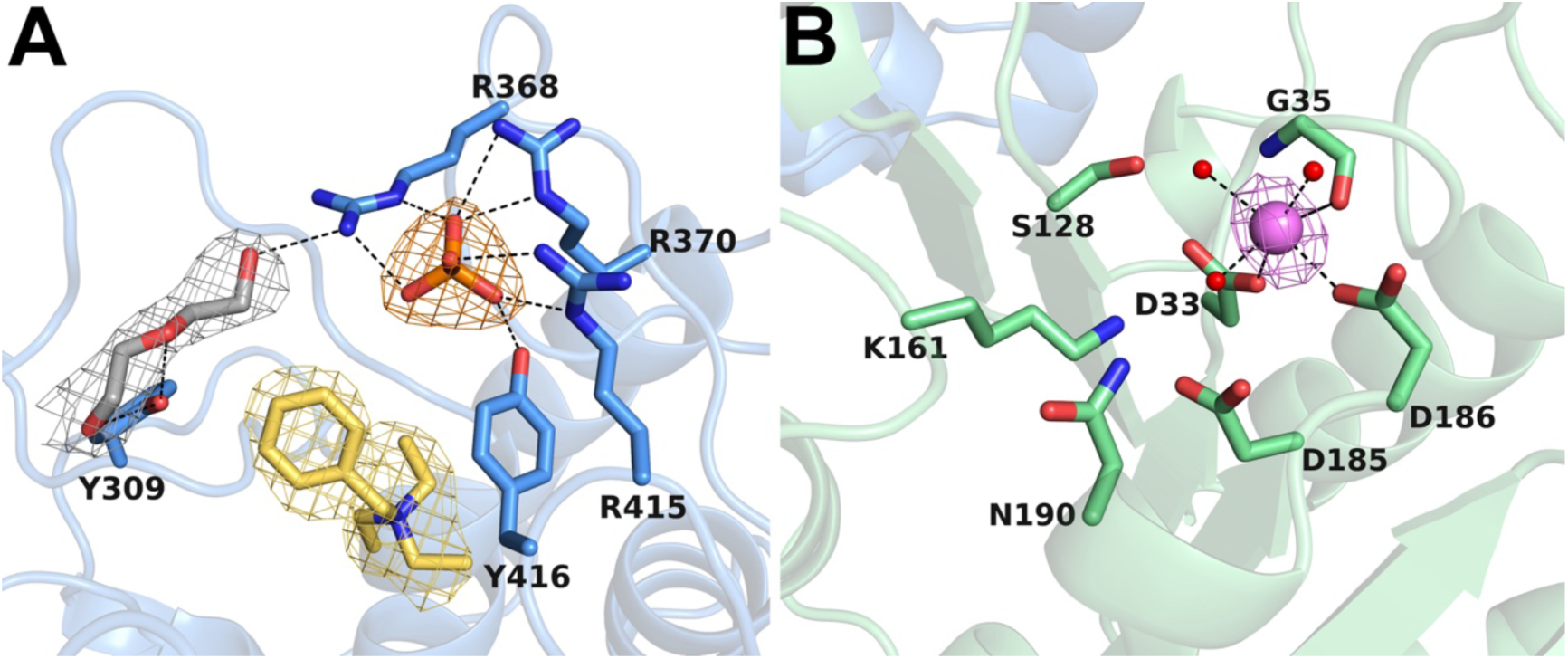
Cyclase domain and phosphatase domain active sites. (A) Cyclase domain in monomer B. Polder omit maps show BTAC (yellow, 4σ contour), P_i_ (orange, 5σ contour), and polyethylene glycol (PEG, grey, 5σ contour) bound in the active site. Hydrogen bonds are indicated by dotted lines. (B) Phosphatase domain in monomer A. The Polder omit map (5σ contour) shows a Mg^2+^ ion (purple) coordinated by D33, D186, the backbone carbonyl of G35, and three water molecules (red spheres) with octahedral geometry.

The HAD-like phosphatase domain of AsDMS contains an α/β Rossmann-like fold in a central core domain. The central β-sheet of the core domain consists of five β-strands (β1–5) in the order 54123 with all βαβ connections being right-handed. HAD-like phosphatases employ a four-loop scaffold within the core domain that positions four key motifs in the active site.^46,47^ The first is the DXD motif, which has diverged to D33-L34-G35 in AsDMS. The first aspartate residue in this motif is responsible for nucleophilic attack at the substrate phosphate group. The second motif is a highly conserved serine or threonine at the end of β2, which corresponds to S128 in AsDMS, and the third motif is a lysine, K161, at the end of the helix between β1 and β4. The second and third motifs stabilize reaction intermediates, with the positive charge of the lysine likely stabilizing the negatively charged phosphate leaving group. The fourth motif at the end of β4 includes acidic residues comprising either a DD, GDxxxD, or GDxxxxD motif; in AsDMS, the DD motif is D185-D186.

HAD-like phosphatases typically exhibit Mg^2+^-dependent catalytic activity,^34,35^ and catalysis by AsDMS accordingly requires Mg^2+^ as mentioned in the previous section. The crystal structure reveals that a single Mg^2+^ ion binds in the phosphatase active site and is octahedrally coordinated by D33, D186, the backbone carbonyl of G35, and three water molecules (Figure 3B). This coordination geometry is consistent with previous observations in other HAD-like phosphatases that the carboxylate group of the first aspartate and the backbone carbonyl of the second aspartate in the DXD motif (which appears as DXG in AsDMS), coordinate to the catalytic Mg^2+^ ion, as well as one aspartate from the fourth motif.^48^

In some HAD-like phosphatases, this core domain is sufficient for activity. However, most also possess cap domains that confer substrate specificity and mediate solvent access to the active site.^49,50^ HAD-like phosphatases can be differentiated based on their cap domains classified as C0, C1, or C2. The AsDMS phosphatase has a C1 cap domain consisting of a ∼70- residue four-helix bundle.

The structure of AsDMS cocrystallized with FSPP was determined at 2.6 Å resolution, revealing that FSPP had been dephosphorylated during cocrystallization to yield one molecule of farnesyl thiol bound in the cyclase active site (Figure 2B, Figure 4). The terminal isoprenoid moiety faces the catalytic DXDT motif, poised as FPP would be for protonation and initiation of the cyclization cascade. However, the partly-extended conformation of the linear isoprenoid is not consistent with that required for substrate binding, i.e., the overall conformation of farnesyl thiol does not reflect the productive FPP conformation that would lead to drimenyl diphosphate generation. However, the isoprenoid conformation would be consistent with the formation of a monocyclic sesquiterpene thiol such as the prematurely quenched product proposed in Figure S4C. As noted above, FSPP is also utilized as a substrate by the cyclase domain over the time course of cocrystallization to generate drimenyl thiol, despite which there is no evidence of drimenyl thiol binding or drimenyl thiodiphosphate binding in either catalytic domain.

**Figure 4.**
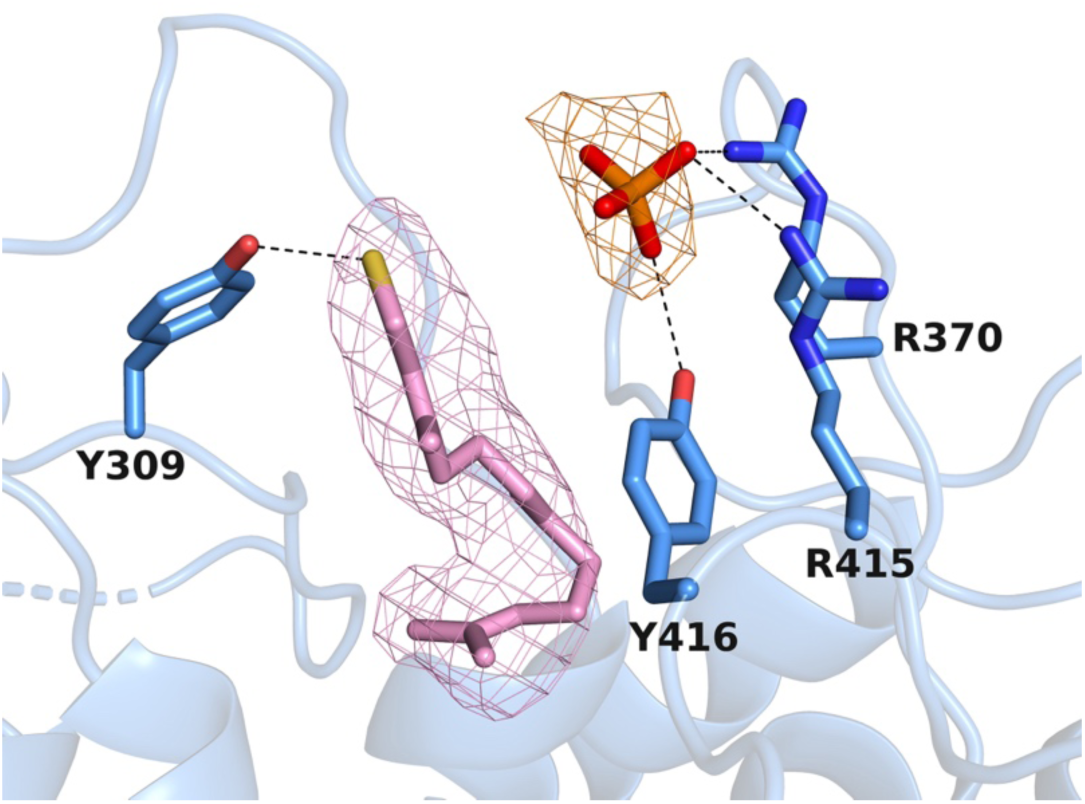
Active site of the cyclase domain in the AsDMS-farnesyl thiol complex (monomer B). The cyclase domain is colored blue. Polder omit maps of farnesyl thiol (pink) and P_i_ (orange) are contoured at 5 and 8.5 σ, respectively. Hydrogen bonds are indicated by dashed lines.

## Discussion

It is instructive to compare the structure of AsDMS with that of the recently reported drimenyl diphosphate synthase from *Streptomyces showdoensis* (SsDMS).^33^ Both AsDMS and SsDMS contain a class II cyclase domain where FPP is converted into drimenyl diphosphate, but only AsDMS contains a phosphatase domain that converts drimenyl diphosphate into drimenol. Accordingly, these enzymes exhibit substantially different domain architectures. SsDMS exhibits the βγ domain architecture characteristic of most class II terpene cyclases, with its active site located at the interdomain interface. In contrast, the class II cyclase domain of AsDMS consists of only a β domain. Comparison of the β domains of AsDMS and SsDMS using DALI^43^ reveals that these two proteins are related by 15% sequence identity, a Z score of 22.3, and an rmsd of 3.5 Å for 493 residues. It is notable that the same cyclization reaction is catalyzed in β domain active sites with such contrasting structures; the lack of a γ domain in AsDMS results in longer loop segments at the top of the active site to protect the binding pocket, much as the γ domain serves this function in SsDMS (Figure 5A).

**Figure 5.**
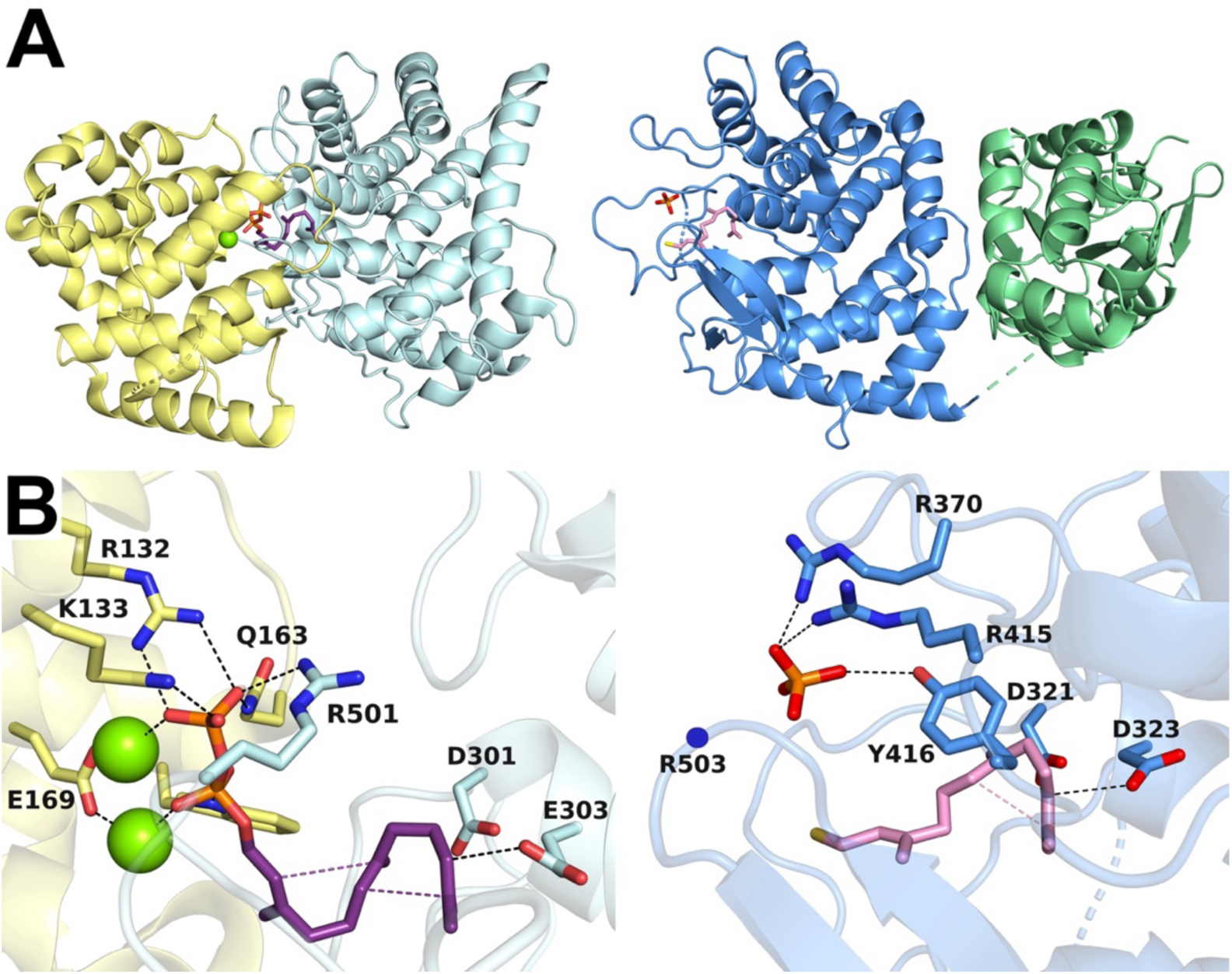
(A) Structural comparison of the D303E SsDMS-FPP complex (left, PDB 7XR7) and the AsDMS-farnesyl thiol complex (right). In SsDMS, the γ domain (yellow) largely caps the active site in the β domain (pale cyan). In AsDMS, longer loop segments serve a similar role in capping the active site in the β domain (blue); the phosphatase domain is green. (B) Close-up views of the active sites in D303E SsDMS (left) and AsDMS (right). Apart from R501, all SsDMS residues involved in Mg^2+^ binding and substrate hydrogen bonding are located in the γ domain. In AsDMS, R370, R415, and Y416 form hydrogen bonds with a P_i_ anion; R503 (blue circle) is disordered in the crystal structure but may become ordered to form a hydrogen bond with the substrate diphosphate group. Dashed black lines indicate hydrogen bonds, and dashed purple and mauve lines indicate trajectories of carbon-carbon bond formation in SsDMS and AsDMS, respectively. A dashed line between the catalytic general acid (D303E in SsDMS, D323 in AsDMS) and the substrate indicates the protonation of the isoprenoid C=C bond that initiates the cyclization cascade. Notably, electrophilic activation of the substrate diphosphate group in AsDMS is achieved solely by hydrogen bonding and not by metal ion coordination.

Cocrystallization of a variant of SsDMS in which the catalytic aspartic acid was replaced by a glutamic acid with FPP and Mg^2+^ enabled the structure determination of an intact enzyme-substrate complex (Figure 5B).^33^ Interestingly, the binding mode of FPP exhibits similarities to that of farnesyl thiol in its complex with AsDMS (Figure 5B). In both complexes, the hydrophobic tail of the isoprenoid ligand is nestled within the hydrophobic pocket of the cyclase. The conformation of FPP is compressed in the SsDMS active site is consistent with the two carbon-carbon bond forming reactions in the cyclization cascade, whereas the conformation of farnesyl thiol is more extended in the AsDMS active site, though it may be poised to enable the first cyclization reaction. Of note, however, the *re* face of the terminal C=C bond is oriented toward the catalytic general acid (D323) in AsDMS, whereas the *si* face of the terminal C=C bond is oriented toward the catalytic general acid (D303E) in SsDMS.

It is also interesting to compare phosphate/diphosphate binding sites between AsDMS and SsDMS. In the AsDMS-BTAC-P_i_ complex, the free phosphate group is stabilized by hydrogen bonds with R368, R370, R415, and Y416, all located on two loops in the β domain (Figure 3A). No other basic residues or metal-binding aspartate or glutamate residues are found in this region. In contrast, in the D303E SsDMS-FPP complex, the diphosphate group of FPP forms hydrogen bonds with R501 in the β domain and R132 and K133 in the γ domain, and is additionally stabilized by coordination to two Mg^2+^ ions coordinated by E169 in the γ domain. None of these residues are conserved in AsDMS, so it appears that Mg^2+^ is not required for FPP binding and activation. Instead, four arginine residues and a tyrosine residue likely form hydrogen bonds with the substrate diphosphate group, based on hydrogen bonds with the phosphate anion observed in AsDMS structures (R368, R370, R415, and Y416; R503 is likely involved as well, but this residue is on a disordered loop at the top of the active site that probably becomes ordered upon substrate binding).

All structural and enzymological data presented above can be integrated into a chemical mechanism for the AsDMS-catalyzed conversion of FPP into drimenol (Figure 6). Upon FPP binding to the class II cyclase domain, the terminal isoprenoid C=C bond is protonated by general acid D323, which triggers concerted C–C bond formation leading to the trans-bicyclic ring system containing a tertiary carbocation. After proton elimination, product drimenyl diphosphate freely diffuses to the phosphatase domain, where it undergoes stepwise dephosphorylation reactions to yield drimenol. Notably, FSPP can undergo the same reaction sequence to generate drimenyl thiol (Figure S4A), but the reaction sequence with this non-native substrate is subject to premature termination based on the generation of farnesyl thiol (Figure S4B) and a putative monocyclic sesquiterpene (Figure S4C). The partly-extended conformation of farnesyl thiol in the cyclase active site, in which only one 6-membered ring appears to be poised for C–C bond formation, could represent the FSPP conformation leading to the putative monocyclic sesquiterpene shown in Figure S4C.

**Figure 6.**
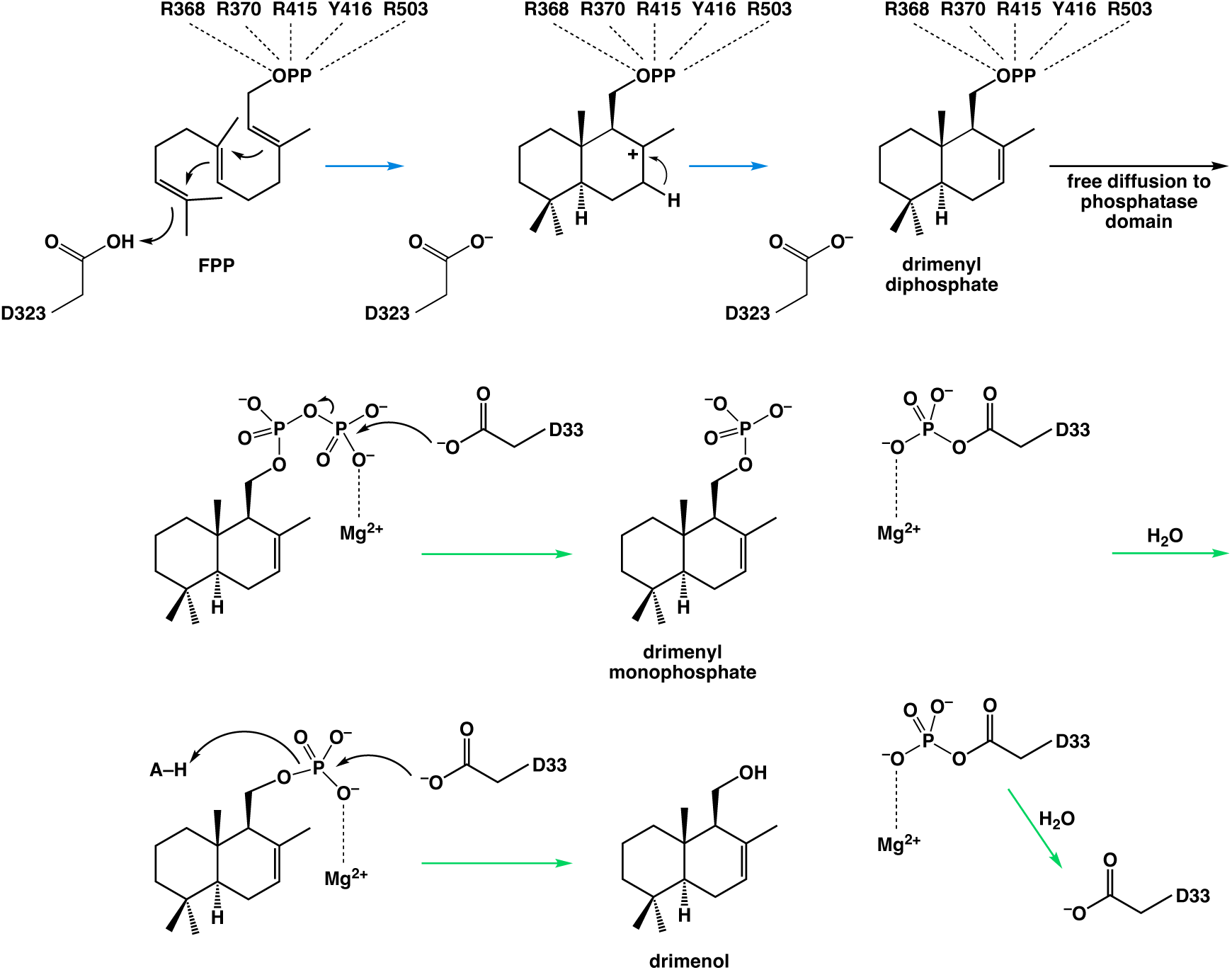
Proposed mechanism for the tandem cyclization-phosphatase reactions catalyzed by AsDMS. FPP is converted to drimenyl diphosphate in the class II cyclase domain (blue arrows), after which drimenyl diphosphate undergoes stepwise dephosphorylation reactions in the phosphatase domain (green arrows). Since the characteristic DXD motif of HAD-like phosphatases appears as DXG in AsDMS, which eliminates the general acid in the phosphatase reaction, the general acid is indicated as A–H above.

## Concluding Remarks

The X-ray crystal structures of AsDMS and its complex with farnesyl thiol are the first of a bifunctional terpene cyclase-phosphatase. Two distinct active sites are observed for the isoprenoid cyclization and dephosphorylation reactions, and enzymological studies show that the dephosphorylation of intermediate drimenyl diphosphate occurs in stepwise fashion to general drimenol. The crystal structure of AsDMS corrects a previous proposal of a single active site for both catalytic activities of drimenol synthase based on molecular modeling studies.^27^ Additionally, the enzymological data reported herein correct a previously proposed catalytic mechanism proceeding through a highly unstable primary carbocation intermediate generated from drimenyl diphosphate, with quenching by a water molecule from bulk solvent to yield the hydroxyl group of drimenol.^27^ Instead, the hydroxyl oxygen of drimenol derives from the prenyl oxygen atom of substrate FPP.

Surprisingly, incubation of AsDMS with FSPP yields drimenyl thiol, in which the thiol sulfur derives from the prenyl sulfur atom of FSPP. Farnesyl thiol and a putative monocyclic sesquiterpene thiol are also generated from FSPP and represent derailment products from the normal isoprenoid cyclization cascade. We anticipate that future studies of AsDMS and its site-specific mutants will set the foundation for engineering “designer cyclases” capable of generating new terpenoids from FPP, FSPP, and perhaps other isoprenoid substrates.

## Methods

### Expression and Purification

A model of AsDMS generated with AlphaFold^51^ revealed that the first 10 residues at the N-terminus had a low predicted local distance difference test (pLDDT) score, which reflected low confidence in the model prediction. Anticipating that these residues might be disordered in the full-length protein, a pET-His6-MBP-TEV-AsDMS plasmid lacking these residues was designed (His6 = hexahistidine purification tag, MBP = maltose binding protein, TEV = tobacco etch virus protease), codon-optimized for expression in *Escherichia coli*, and purchased from GenScript (Piscataway, NJ) (Scheme 1).

**Scheme 1.**
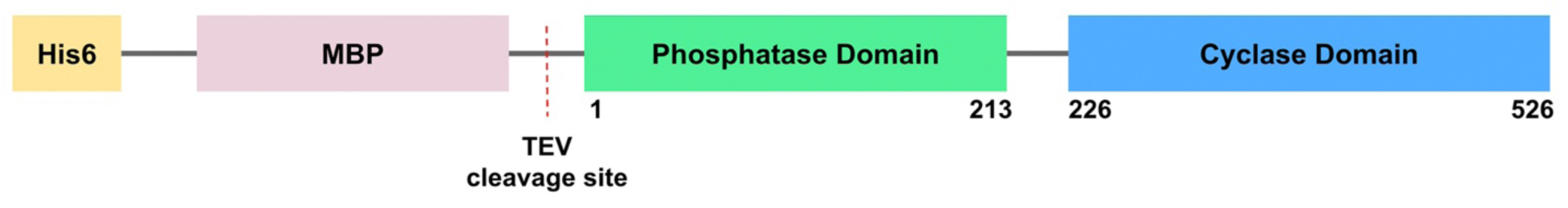
Expression plasmid for AsDMS truncation variant

The AsDMS plasmid was transformed into BL21(DE3) *E. coli* cells (New England Biolabs) and grown overnight at 37° C on Luria–Bertani (LB)-agar plates containing 50 µg/mL kanamycin. Single colonies were picked and used to inoculate 250 mL of LB media supplemented with 50 µg/mL kanamycin. This 250-mL starter culture was incubated overnight in an orbital shaker (175 rpm, 37° C) and subsequently used to inoculate 12 x 1-L flasks containing 1-L of LB media supplemented with 50 µg/mL kanamycin. The 1-L cultures were incubated (175 rpm, 37° C) until each reached an optical density at 600 nm (OD_600_) of ∼0.6-1.0. Protein expression was then induced by the addition of isopropyl-β-D-thiogalactopyranoside (IPTG) to a final concentration of 0.5 mM, and cultures were incubated overnight in the orbital shaker (175 rpm, 18° C) for expression. Cells were pelleted via centrifugation (20 min, 6000 rpm, 4° C), and the pellet was stored at −80° C until purification.

Purification of AsDMS began with resuspension of the cell pellet in buffer A [50 mM sodium phosphate (monobasic, monohydrate, pH 7.3), 500 mM NaCl, and 20% glycerol]. The suspended pellet was treated with 100 mg lysozyme (GoldBio), 8 mg DNase I – Grade II (Roche Diagnostics), and one EDTA-free cOmplete Mini Protease Inhibitor Tablet (Roche Diagnostics) and stirred at room temperature for 1 h. The cell suspension was then lysed using a Q700 sonicator (Qsonica) at an amplitude of 30% for 10 min (cycling between 1 s on/2 s off). Cell lysate was clarified by centrifugation at 18,000 rpm for 1 h at 4° C.

Supernatant was loaded at 3 mL/min onto a 5-mL HiTrap TALON column (GE Healthcare) preequilibrated with buffer A, and eluted via a 50-mL gradient to buffer B [50 mM sodium phosphate (monobasic, monohydrate, pH 7.3), 500 mM NaCl, 200 mM imidazole, 20% glycerol]. Selected fractions were analyzed using sodium dodecyl sulfate-polyacrylamide gel electrophoresis (SDS-PAGE). Fractions determined to contain eluted protein were pooled and treated with 3 mg of TEV protease while undergoing dialysis into buffer A at 4° C.

The dialyzed sample of AsDMS was loaded onto a tandem 5-mL HiTrap TALON/MBP Trap column. The flow-through was analyzed by SDS-PAGE, and fractions containing eluted protein were collected, pooled, and concentrated to approximately 5 mL using a 50-kDa cutoff Amicon Ultra-15 Centrifugal Filter Unit. The sample was then loaded onto a HiLoad 26/60 Superdex 200-pg size-exclusion column (GE Healthcare), which was pre-equilibrated with buffer C [20 mM Tris-HCl (pH 7.3), 150 mM NaCl, 1 mM tris(2-carboxyethyl)phosphine (TCEP), 10% glycerol]. Fractions containing eluted protein were determined to be >95% pure based on SDS-PAGE and were pooled and concentrated to approximately 10–30 mg/mL.

### Enzyme Activity

For identification and quantification of reaction products, activity assays were conducted using gas chromatography-mass spectrometry (GC-MS) with an Agilent 8890 GC/5597C MSD system equipped with a J&W HP-5 ms Ultra Inert GC capillary column (30 m × 0.25 mm × 0.25 μm). Assays were performed in triplicate on a 200-μL scale using an enzyme concentration of 5 μM in reaction buffer [20 mM Tris-HCl (pH 7.3), 150 mM NaCl, 1 mM TCEP, 10% glycerol] with 2 mM MgCl_2_•6H_2_O. Farnesyl diphosphate (FPP) was purchased from Isoprenoids.com and resuspended in 7:3 methanol/10 mM NH_4_HCO_3_ at a concentration of 10 mM. Reactions were initiated by addition of 10 μL of the 10 mM FPP stock followed immediately by carefully layering 200 μL of hexanes containing an internal standard of 125 mM β-pinene over the aqueous phase. Reactions were incubated at room temperature overnight, and were quenched by vortexing for 10 s, followed by centrifugation for 15 s at 12,000 rpm. An aliquot of the organic phase (100 μL) was removed for GC-MS analysis. For all samples, the temperature program began with an oven temperature of 60 °C sustained for 2 min, followed by a ramp at 20°C/min to reach 240 °C. A positive EI mode was used to collect MS data after a solvent delay of 3 min. Product drimenol was characterized through comparison of retention time and mass spectrum with data archived in the National Institute of Standards and Technology (NIST) database.

To test enzymatic activity in the presence of various metals, assays were performed as described above, except the enzyme-buffer mixture with no added metal was preincubated with 2 mM EDTA to chelate any residual metal ions remaining from the purification process. After 30 min, 4 mM of the metal of interest was added and reactions were initiated upon the addition of FPP.

To determine whether a water molecule from bulk solution is the source of the hydroxyl group of drimenol, a 570-μL sample containing 5 μM enzyme in reaction buffer [20 mM Tris-HCl (pH 7.3), 150 mM NaCl, 1 mM TCEP] with 2 mM MgCl_2_•6H_2_O was lyophilized for 2 h. The sample was resolubilized in 570 μL H_2_^18^O and split into three vials, and reactions were initiated upon the addition of 10 μL of the 10 mM FPP stock solution. Samples were incubated overnight and subject to GC-MS analysis, as described above.

For assays using the weakly reactive substrate FSPP, the same approach outlined above was utilized except reactions were initiated upon the addition of 2.5 μL of a 40 mM FSPP stock solution.

To ascertain whether AsDMS catalyzes stepwise phosphate (P_i_) removal rather than removal of intact pyrophosphate (PP_i_) from the substrate, we modified the EnzChek Pyrophosphate Detection Assay Kit (ThermoFisher Scientific) to detected free phosphate in solution.^32^ Purine nucleoside phosphorylase (PNP) converts the substrate 2-amino6-mercapto- 7-methylpurine riboside (MESG) into ribose 1-phosphate and 2-amino6-mercapto-7- methylpurine in the presence of P_i_, which results in an absorbance shift from 330 nm to 360 nm. Typically, inorganic pyrophosphatase in included to break down pyrophosphate into free P_i_; by excluding inorganic pyrophosphatase, the assay will only yield an absorbance shift if P_i_, rather than PP_i_ is produced by AsDMS. All assays were performed in triplicate on a 100-µL scale using 1 µM AsDMS reaction buffer [20 mM Tris-HCl (pH 7.3), 150 mM NaCl, 1 mM TCEP], 50 µM substrate, 0.1 U of PNP, and 200 µM MESG. Absorbance at 360 nm was measured using UV-Vis Spectroscopy after incubation for 1 h at 22°C. A control reaction in the absence of enzyme was used as a baseline. Absorbance was converted to P_i_ concentration by comparison to a standard curve.

### Crystal Structure Determination

Crystallization was achieved by the sitting-drop vapor diffusion method, using a Mosquito pipetting robot (SPT Labtech) to screen crystallization conditions. Crystals were generated using two different conditions:

### Condition 1

A 100-nL drop of protein solution [13 mg/mL AsDMS, 20 mM Tris-HCl (pH 7.3), 150 mM NaCl, 1 mM TCEP, 10% glycerol, 2 mM farnesyl *S*-thiolodiphosphate (FSPP), MgCl_2_·6H_2_O] was added to a 100-nL drop of precipitant solution [0.1 M Sodium citrate tribasic dihydrate pH 5.5, 26% v/v Jeffamine® ED-2001 pH 7.0] and equilibrated against a 50-µL reservoir of precipitant solution.

### Condition 2

A 100-nL drop of protein solution [13 mg/mL AsDMS, 20 mM Tris-HCl (pH 7.3), 150 mM NaCl, 1 mM TCEP, 10% glycerol, 2 mM benzyltriethylammonium chloride (BTAC), 2 mM sodium pyrophosphate, 2 mM MgCl_2_·6H_2_O] was added to a 100-nL drop of precipitant solution [0.1 M 4-(2-hydroxyethyl)piperazine-1-ethane-sulfonic acid (HEPES) (pH 7.5), 0.2 M sodium chloride, 25% w/v PEG 3350] and equilibrated against a 50-µL reservoir of precipitant solution.

For each crystallization condition, crystals appeared within 14 days and were cryoprotected in mother liquor augmented with 20% (v/v) glycerol prior to flash-cooling in liquid nitrogen.

All X-ray diffraction data were collected at the 17-ID-1 AMX beamline at National Synchrotron Light Source II (NSLS-II) at Brookhaven National Laboratory (Upton, NY). Each data set was indexed, integrated, and scaled with XDS^52^ and merged with AIMLESS as part of the autoPROC pipeline.^53^ The data set from condition 1 was phased using molecular replacement with the PHASER module of PHENIX,^54,55^ and an AlphaFold-predicted^51^ structure of AsDMS was used as the search model. The space group was authenticated using Zanuda,^56^ and the output model underwent a round of manual model building using Coot.^57^ Iterative rounds of manual model building and crystallographic refinement were performed using Coot and PHENIX. During model building, side chains were deleted which exhibited no or poor electron density in the 2|F|_o_ – |F|_c_ map contoured at 1.5σ. The data set from condition 2 was phased using the refined crystal structure from condition 1 as the search model. The structure from condition 2 was refined in the same manner as the structure from condition 1. Bound ligands and solvent molecules, including buffer components (glycerol and PEG fragments) were added to the model in the final stages of refinement. The final refined structure was validated using MolProbity.^58^ All data collection and refinement statistics are recorded in Table 1.

**Table 1.** Data collection and refinement statistics.

| Complex | AsDMS-BTAC-P<sub>i</sub>-Mg<sup>2+</sup> | AsDMS-farnesyl thiol-P<sub>i</sub> |
| --- | --- | --- |
| Space group | C2 | *P*4<sub>1</sub>2<sub>1</sub>2 |
| a,b,c (Å) | 179.90, 76.25, 106.43 | 154.74, 154.74, 132.98 |
| α, β, γ (°) | 90.00, 111.68, 90.00 | 90.00, 90.00, 90.00 |
| R<sub>merge</sub><sup>b</sup> | 0.174 (1.297) | 0.406 (3.808) |
| R<sub>pim</sub><sup>c</sup> | 0.095 (0.698) | 0.101 (0.928) |
| CC<sub>1/2</sub><sup>d</sup> | 0.989(0.330) | 0.993 (0.339) |
| Redundancy | 4.5(4.6) | 17.0 (17.6) |
| Completeness (%) | 99.9 (99.1) | 99.9 (99.4) |
| I/σ | 6.4 (1.3) | 7.0 (0.9) |
| Refinement |  |  |
| Resolution (Å) | 2.23 | 2.64 |
| No. reflections | 65482 | 47665 |
| R<sub>work</sub>/R<sub>free</sub><sup>e</sup> | 0.206/0.249<br>(0.293)/(0.347) | 0.217/0.265<br>(0.301)/(0.345) |
| Number of atoms<sup>f</sup> |  |  |
| Protein | 7606 | 7552 |
| Ligands | 77 | 68 |
| Solvent | 197 | 94 |
| Average B factors (Å<sup>2</sup>) |  |  |
| Protein | 34 | 50 |
| Ligands | 43 | 60 |
| Solvent | 36 | 45 |
| RMS deviations |  |  |
| Bond lengths (Å) | 0.007 | 0.008 |
| Bond angles (°) | 0.8 | 0.9 |
| Ramachandran plot<sup>g</sup> |  |  |
| Favored (%) | 98.35 | 94.98 |
| Allowed (%) | 1.65 | 4.71 |
| Outliers (%) | 0.00 | 0.31 |
| Molprobity score | 1.38 | 2.07 |
| PDB Entry | 9MHS | 9MLO |

<sup>a</sup>Values in parentheses refer to the highest-resolution shell of data.
<sup>b</sup>R<sub>merge</sub> = $\sum_h \sum_i |I_{i,h} - \langle I \rangle_h| / \sum_h \sum_i I_{i,h}$ , where $\langle I \rangle_h$ is the average intensity calculated for reflection *h* from *i* replicate measurements.
<sup>c</sup>R<sub>p.i.m.</sub> = $(\sum_h (1/(N-1))^{1/2} \sum_i |I_{i,h} - \langle I \rangle_h|) / \sum_h \sum_i I_{i,h}$ , where *N* is the number of reflections and $\langle I \rangle_h$ is the average intensity calculated for reflection *h* from replicate measurements.
<sup>d</sup>Pearson correlation coefficient between random half-datasets.
<sup>e</sup>R<sub>work</sub> = $\sum ||F_o| - |F_c|| / \sum |F_o|$ for reflections contained in the working set. $|F_o|$ and $|F_c|$ are the observed and calculated structure factor amplitudes, respectively. R<sub>free</sub> is calculated using the same expression for reflections contained in the test set held aside during refinement.
<sup>f</sup>Per asymmetric unit.
<sup>g</sup>Calculated with MolProbity.

In the final refined crystal structure from condition 1, some segments of AsDMS are disordered and hence excluded from the final model. In monomer A, these segments include 13 residues at the N-terminus (N1–L13), the interdomain linker (I214–M225), and a small loop in the cyclase domain (N297–Q301). In monomer B, these segments include an N-terminal segment (N1–Q18), the interdomain linker (I214–M225), and a small loop in the cyclase domain (H298–Q301).

In the final refined crystal structure from condition 2, disordered segments are also observed in both monomers and thus excluded from the final model. In monomer A, these segments include ten residues at the N-terminus (N1–T10) a loop in the phosphatase domain (F46–T50), and the interdomain linker (G216–M225). In monomer B, these segments include the N-terminus (N1–L20), a loop in the phosphatase domain (W41–D54), and the interdomain linker (N205–K206 and G216–M225).

## Data Availability

The atomic coordinates and crystallographic structure factors of the AsDMS-BTAC-P_i_-Mg^2+^ complex and the AsDMS-farnesyl thiol-P_i_ complex have been deposited in the Protein Data Bank (www.rcsb.org) with accession codes 9MHS and 9MLO, respectively.

## Funding

This research was supported by NIH grant GM56838 to D.W.C. M.N.G. was supported by Chemistry-Biology Interface NIH Training Grant T32 GM133398. K.R.O. was supported by the Vagelos Program in Molecular Life Sciences at the University of Pennsylvania.

## Conflict of Interest Statement

The authors declare no competing interests.

## ACKNOWLEDGMENTS

We are especially grateful to Dr. Samuel Eaton for his scientific guidance and mentorship during the early stages of this work. We also thank Drs. Juana Goulart Stollmaier and Kollin Schultz for helpful scientific discussions and assistance with experimental techniques. This work is based on research conducted at beamline 17-ID-1 (AMX) of the National Synchrotron Light Source II, a DOE Office of Science User Facility operated for the DOE Office of Science by Brookhaven National Laboratory under Contract DE-SC0012704. The Center for BioMolecular Structure (CBMS) is primarily supported by the National Institutes of Health, NIGMS, through a Center Core P30 Grant (P30GM133893) and by the DOE Office of Biological and Environmental Research (KP1605010).

## Notes

### Competing Interest Statement

The authors have declared no competing interest.

## References

1. Tholl, D. (2006) Terpene synthases and the regulation, diversity and biological roles of terpene metabolism. Curr. Opin. Plant Biol. 9, 297–304.

2. Cane, D. E., Ikeda, H. (2012) Exploration and mining of the bacterial terpenome. Acc. Chem. Res. 45, 463–472.

3. Quin, M. B., Flynn, C. M., Schmidt-Dannert, C. (2014) Traversing the fungal terpenome. Nat. Prod. Rep. 31, 1449–1473.

4. Walsh, C. T., Tang, Y. (2017) Natural product biosynthesis: chemical logic and enzymatic machinery. Royal Society of Chemistry: Croydon, U.K., 794 pp.

5. Tetali, S. D. (2017) Terpenes and isoprenoids: a wealth of compounds for global use. Planta 249, 1–8.

6. Fan, M., Yuan, S., Li, L., Zheng, J., Zhao, D., Wang, C., Wang, H., Liu, X., Liu, J. (2023) Application of terpenoid compounds in food and pharmaceutical products. Fermentation 9, 119.

7. Buckingham, J. (1993). Dictionary of Natural Products. Chapman & Hall: London, 8584 pp.

8. https://dnp.chemnetbase.com.

9. Poulter, C. D., Rilling, H. C. (1978) The prenyl transfer reaction. Enzymatic and mechanistic studies of the 1’-4 coupling reaction in the terpene biosynthetic pathway. Acc. Chem. Res. 11, 307–313.

10. Kellogg, B. A., Poulter, C. D. (1997) Chain elongation in the isoprenoid biosynthetic pathway. Curr. Opin. Chem. Biol. 1, 570–578.

11. Cane, D. E. (1990). Enzymatic formation of sesquiterpenes. Chem. Rev. 90, 1089–1103.

12. Christianson, D. W. (2006) Structural biology and chemistry of the terpenoid cyclases. Chem. Rev. 106, 3412–3442.

13. Degenhardt, J., Köllner, T. G., Gershenzon, J. (2009) Monoterpene and sesquiterpene synthases and the origin of terpene skeletal diversity in plants. Phytochemistry 70, 1621–1637.

14. Christianson, D. W. (2017) Structural and chemical biology of terpenoid cyclases. Chem. Rev. 117, 11570–11648.

15. Whitehead, J. N., Leferink, N. G. H., Johannissen, L. O., Hay, S., Scrutton, N. S. (2023) Decoding catalysis by terpene synthases. ACS Catal. 13, 12774–12802.

16. Oldfield, E., Lin, F.-Y. (2011) Terpene biosynthesis: modularity rules. Angew. Chem. Int. Ed. 51, 1124–1137.

17. Chen, M., Harris, G. G., Pemberton, T. A., Christianson, D. W. (2016) Multi-domain terpenoid cyclase architecture and prospects for proximity in bifunctional catalysis. Curr. Opin. Struct. Biol. 41, 27–37.

18. Faylo, J. L., Ronnebaum, T. A., Christianson, D. W. (2021) Assembly-line catalysis in bifunctional terpene synthases. Acc. Chem. Res. 54, 3780–3791.

19. Vogel, B. S., Wildung, M. R., Vogel, G., Croteau, R. (1996) Abietadiene synthase from grand fir (*Abies grandis*). cDNA isolation, characterization, and bacterial expression of a bifunctional diterpene cyclase involved in resin acid biosynthesis. J. Biol. Chem. 271, 23262–23268.

20. Peters, R. J., Ravn, M. M., Coates, R. M., Croteau, R. B. (2001) Bifunctional abietadiene synthase: free diffusive transfer of the (+)-copalyl diphosphate intermediate between two distinct active sites. J. Am. Chem. Soc. 123, 8974–8978.

21. Zhou, K., Gao, Y., Hoy, J. A., Mann, F. M., Honzatko, R. B., Peters, R. J. (2012) Insights into diterpene cyclization from structure of bifunctional abietadiene synthase from *Abies grandis*. J. Biol. Chem. 287, 6840–6850.

22. Toyomasu, T., Tsukahara, M., Kaneko, A., Niida, R., Mitsuhashi, W., Dairi, T., Kato, N., Sassa, T. (2007) Fusicoccins are biosynthesized by an unusual chimera diterpene synthase in fungi. Proc. Natl. Acad. Sci. USA 104, 3084–3088.

23. Pemberton, T. A., Chen, M., Harris, G. G., Chou, W. K. W., Duan, L., Köksal, M., Genshaft, A. S., Cane, D. E., Christianson, D. W. (2017) Exploring the influence of domain architecture on the catalytic function of diterpene synthases. Biochemistry 56, 2010–2023.

23. Chen, M., Chou, W. K. W., Toyomasu, T., Cane, D. E., Christianson, D. W. (2016) Structure and function of fusicoccadiene synthase, a hexameric bifunctional diterpene synthase. ACS Chem. Biol. 11, 889–899.

24. Faylo, J. L., van Eeuwen, T., Kim, H. J., Gorbea Colón, J. J., Garcia, B. A., Murakami, K., Christianson, D. W. (2021) Structural insight on assembly-line catalysis in terpene biosynthesis. Nat. Commun. 12, 3487.

26. Faylo, J. L., van Eeuwen, T., Gupta, K., Murakami, K., Christianson, D. W. (2022) Transient prenyltransferase–cyclase association in fusicoccadiene synthase, an assembly-line terpene synthase. Biochemistry 61, 2417–2430.

26. Wenger, E. S., Schultz, K., Marmorstein, R., Christianson, D. W. (2024) Engineering substrate channeling in a bifunctional terpene synthase. Proc. Natl. Acad. Sci. USA 121, e2408064121.

27. Vo, N. N. Q., Nomura, Y., Kinugasa, K., Takagi, H., Takahashi, S. (2022) Identification and characterization of bifunctional drimenol synthases of marine bacterial origin. ACS Chem. Biol. 17, 1226–1238.

28. Montenegro, I. J.; del Corral, S.; Diaz Napal, G. N.; Carpinella, M. C.; Mellado, M.; Madrid, A. M.; Villena, J.; Palacios, S. M.; Cuellar, M. A. (2018). Antifeedant effect of polygodial and drimenol derivatives against *Spodoptera frugiperda* and *Epilachna paenulata* and quantitative structure-activity analysis. Pest Manag. Sci. 74, 1623–1629.

29. Yang, Z.; Chan, K. W.; Bakar, M. Z. A.; Deng, X. (2024). Unveiling drimenol: a phytochemical with multifaceted bioactivities. Plants 13, 2492.

30. Gomez, G. A., Morisseau, C., Hammock, B. D., Christianson, D. W. (2004) Structure of human epoxide hydrolase reveals mechanistic inferences on bifunctional catalysis in epoxide and phosphate ester hydrolysis. Biochemistry 43, 4716–4723.

31. Enayetallah, A. E., Grant, D. F. (2006) Effects of human soluble epoxide hydrolase polymorphisms on isoprenoid phosphate hydrolysis. Biochem. Biophys. Res. Commun. 341, 254–260.

32. Lloyd, A. J., Thomann, H. U., Ibba, M., Söll, D. (1995) A broadly applicable continuous spectrophotometric assay for measuring aminoacyl-tRNA synthetase activity. Nucleic Acids Res. 23, 2886–2892.

33. Pan, X., Du, W., Zhang, X., Lin, X., Li, F.-R., Yang, Q., Wang, H., Rudolf, J. D., Zhang, B., Dong, L.-B. (2022) Discovery, structure, and mechanism of a class II sesquiterpene cyclase. J. Am. Chem. Soc. 144, 22067–22074.

34. Allen, K. N., Dunaway-Mariano, D. (2004) Phosphoryl group transfer: evolution of a catalytic scaffold. Trends Biochem. Sci. 29, 495–503.

35. Allen, K. N., Dunaway-Mariano, D. (2016) Catalytic scaffolds for phosphoryl group transfer. Curr. Opin. Struct. Biol. 41, 172–179.

36. Krissinel, E., Henrick, K. (2007) Inference of macromolecular assemblies from crystalline state. J. Mol. Biol. 372, 774–797.

37. Pan, X., Rudolf, J. D., Dong, L. (2024) Class II terpene cyclases: structures, mechanisms, and engineering. Natural Product Reports, 41(3), 42–433.

38. Wendt, K. U., Schulz, G. E. (1998) Isoprenoid biosynthesis: manifold chemistry catalyzed by similar enzymes. Structure 6, 127–133.

39. Wendt, K. U., Poralla, K., Schulz, G. E. (1997) Structure and function of a squalene cyclase. Science 277, 1811–1815.

40. Köksal, M., Hu, H., Coates, R. M., Peters, R. J., Christianson, D. W. (2011) Structure and mechanism of the diterpene cyclase *ent*-copalyl diphosphate synthase. Nat. Chem. Biol. 7, 431–433.

41. Köksal, M., Potter, K., Peters, R. J., Christianson, D. W. (2014) 1.55 Å-resolution structure of *ent*-copalyl diphosphate synthase and exploration of general acid function by site-directed mutagenesis. Biochim. Biophys. Acta 1840, 184–190.

42. Moosmann, P., Ecker, F., Leopold-Messer, S., Cahn, J. K. B., Dieterich, C. L., Groll, M., Piel, J. (2020) A monodomain class II terpene cyclase assembles complex isoprenoid scaffolds. Nat. Chem. 12, 968–972.

43. Holm, L. (2022) Dali server: structural unification of protein families. Nucleic Acids Res. 50, W210–W215.

44. Alexander, R. S.; Kanyo, Z. F.; Chirlian, L. E.; Christianson, D. W. (1990) Stereochemistry of phosphate-Lewis acid interactions: implications for nucleic acid structure and recognition. J. Am. Chem. Soc. 112, 933–937.

45. Aaron, J. A., Lin, X., Cane, D. E., Christianson, D. W. (2010) Structure of *epi*-isozizaene synthase from *Streptomyces coelicolor* A3(2), a platform for new terpenoid cyclization templates. Biochemistry 49, 1787–1797.

46. Allen K. N., Dunaway-Mariano D. (2009) Markers of fitness in a successful enzyme superfamily. Curr. Opin. Struct. Biol. 19, 658–665.

47. Burroughs, A. M., Allen, K. N., Dunaway-Mariano, D., Aravind, L. (2006) Evolutionary genomics of the HAD superfamily: understanding the structural adaptations and catalytic diversity in a superfamily of phosphoesterases and allied enzymes. J. Mol. Biol. 361, 1003–1034.

48. Kuznetsova, E., Nocek, B., Brown, G., Makarova, K. S., Flick, R., Wolf, Y. I., Khusnutdinova, A., Evdokimova, E., Jin, K., Tan, K., Hanson, A. D., Hasnain, G., Zallot, R., de Crécy-Lagard, V., Babu, M., Savchenko, A., Joachimiak, A., Edwards, A. M., Koonin, E. V., Yakunin, A. F. (2015) Functional diversity of haloacid dehalogenase superfamily phosphatases from *Saccharomyces cerevisiae*: biochemical, structural, and evolutionary insights. J. Biol. Chem. 290, 18678–18698.

49. Huang, H., Pandya, C., Liu, C., Al-Obaidi, N. F., Wang, M., Zheng, L., Keating, S. T., Aono, M., Love, J. D., Evans, B., Seidel, R. D., Hillerich, B. S., Garforth, S. J., Almo, S. C., Mariano, P. S., Dunaway-Mariano, D., Allen, K. N., Farelli, J. D. (2015) Panoramic view of a superfamily of phosphatases through substrate profiling. Proc. Natl. Acad. Sci. USA 112, E1974–E1983.

50. Lahiri, S. D., Zhang, G., Dai, J., Dunaway-Mariano, D., Allen, K. N. (2004) Analysis of the substrate specificity loop of the HAD superfamily cap domain. Biochemistry 43, 2812–2820.

51. Jumper, J., Evans, R., Pritzel, A. et al. (2021) Highly accurate protein structure prediction with AlphaFold. Nature 596, 583–589.

52. Kabsch, W. (2010). XDS. Acta Cryst. D66, 125–132.

53. Evans, P. R., Murshudov, G. N. (2013). How good are my data and what is the resolution? Acta Cryst. D69, 1204–1214.

54. McCoy, A. J., Grosse-Kunstleve, R. W., Adams, P. D., Winn, M. D., Storoni, L. C., Read, R. J. (2007) Phaser crystallographic software. J. Appl. Crystallogr. 40, 658–674.

55. Adams, P. D., Afonine, P. V., Bunkóczi, G., Chen, V. B., Davis, I. W., Echols, N., Headd, J. J., Hung, L. W., Kapral, G. J., Grosse-Kunstleve, R. W., McCoy, A. J., Moriarty, N. W., Oeffner, R., Read, R. J., Richardson, D. C., Richardson, J. S., Terwilliger, T. C., Zwart, P. H. (2010) PHENIX: a comprehensive Python-based system for macromolecular structure solution. Acta Cryst. D66, 213–221.

56. Lebedev, A. A., & Isupov, M. N. (2014). Space-group and origin ambiguity in macromolecular structures with pseudo-symmetry and its treatment with the program Zanuda. Acta Cryst. D70, 2430–2443. 10.1107/S1399004714014795

57. Emsley, P., Lohkamp, B., Scott, W. G., Cowtan, K. (2010) Features and development of Coot. Acta Cryst. D66, 486–501.

58. Chen, V. B., Arendall, W. B., Headd, J. J., Keedy, D. A., Immormino, R. M., Kapral, G. J., Murray, L. W., Richardson, J. S., Richardson, D. C. (2010) MolProbity: all-atom structure validation for macromolecular crystallography. Acta Cryst. D66, 12–21.

