## Supporting Information for "Crystal Structure and Catalytic Mechanism of Drimenol Synthase, a Bifunctional Terpene Cyclase-Phosphatase"

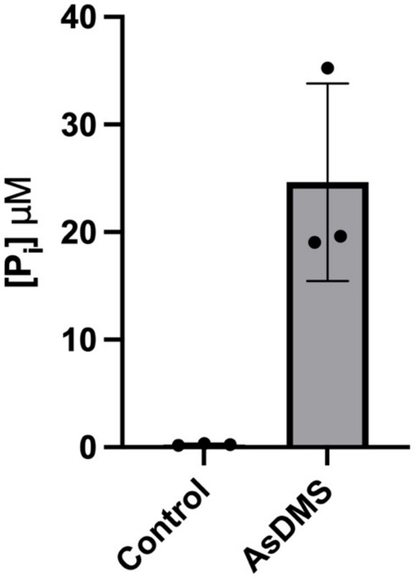


**Figure S1.** EnzChek assay results showing a robust signal for $P_i$ in the AsDMS reaction even though inorganic pyrophosphatase (IPPase) was omitted from the assay. This result indicates that 2 $P_i$ anions instead of one inorganic pyrophosphate anion ($PP_i$) are coproducts of the AsDMS phosphatase reaction. Error bars represent the standard deviation of three independent measurements.

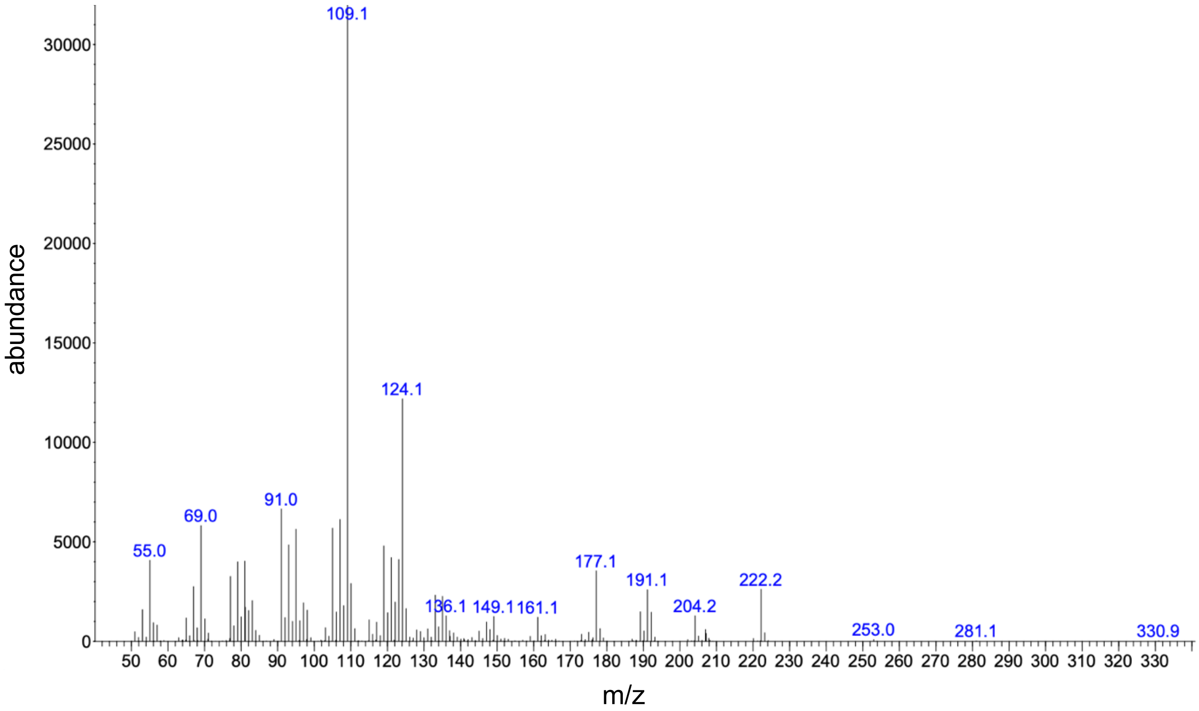


**Figure S2.** Mass spectrum of drimenol product formed when drimenol synthase is incubated with FPP in $H_2{}^{18}O$ buffer. The molecular weight of drimenol remains at 222.2; if $^{18}O$ were incorporated into drimenol, its molecular weight would be 224.2. Small, spurious peaks are observed at m/z = 253.0, 281.1, and 330.9 and correspond to an impurity.

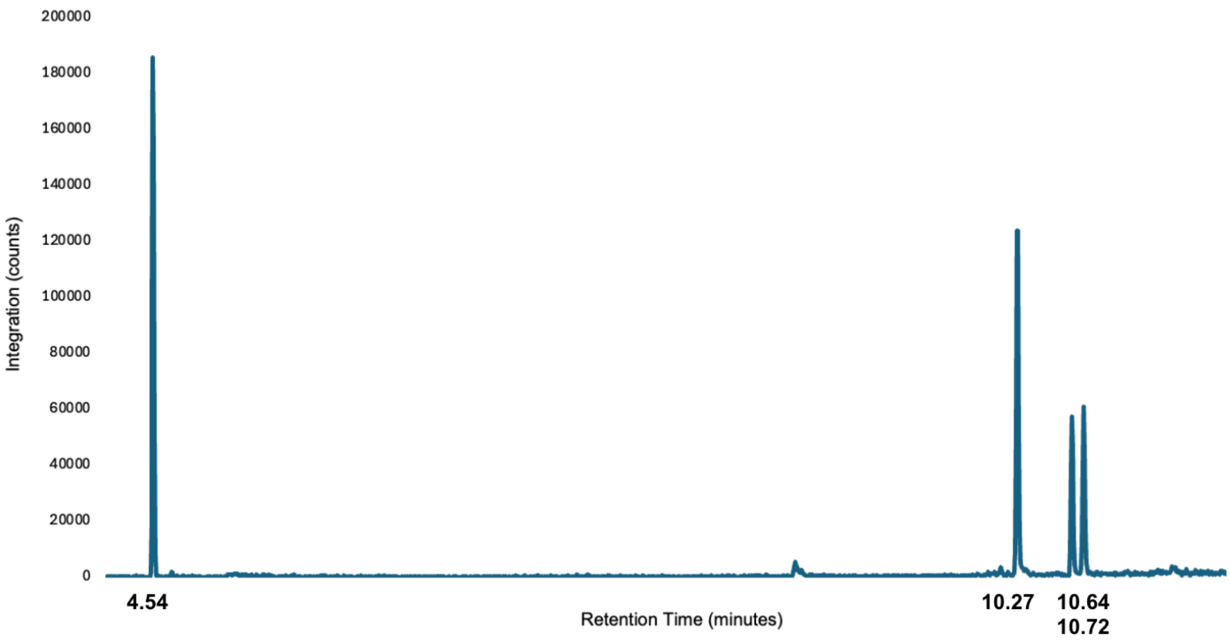


**Figure S3**.Gas chromatogram of products formed when AsDMS is incubated with FSPP. The peak at 4.54 min corresponds to a monoterpene (β-pinene) standard.

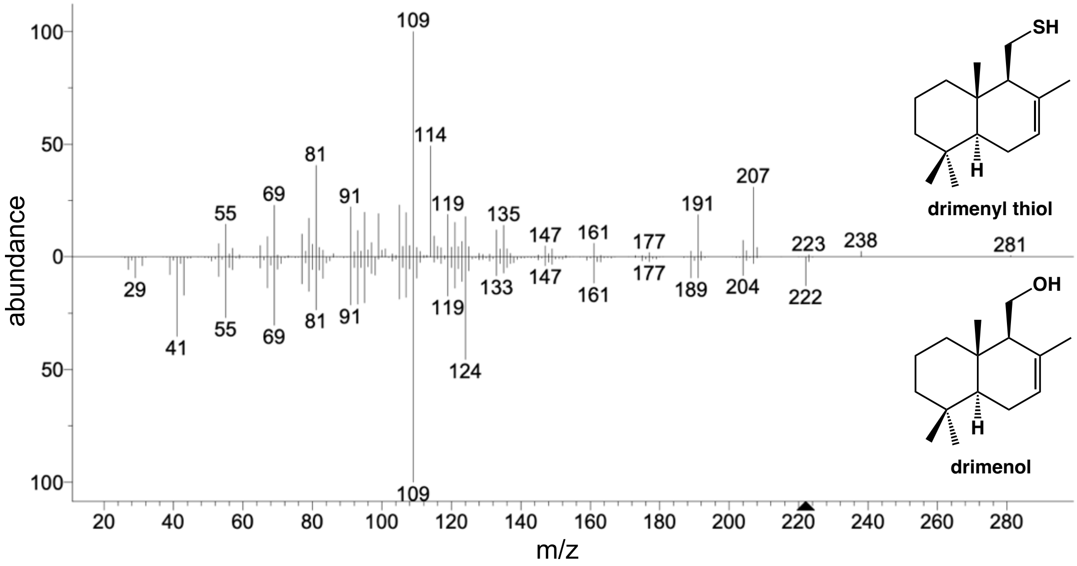


**Figure S4A**. Butterfly plot of mass spectra of drimenol from the NIST database (bottom) and the cyclized product produced when AsDMS is incubated with FSPP (top) showing nearly identical fragmentation patterns. The expected molecular weight of drimenol is 222. The molecular weight of the cyclized product is 238, which indicates incorporation of sulfur (atomic weight: 32) instead of oxygen (atomic weight: 16). A small, spurious peak is observed at m/z = 281 and corresponds to an impurity; this spurious peak is also observed in the mass spectrum of drimenol in Figure S2. The peak at m/z = 207 results from leaching of the polystyrene stationary phase of the GC column.

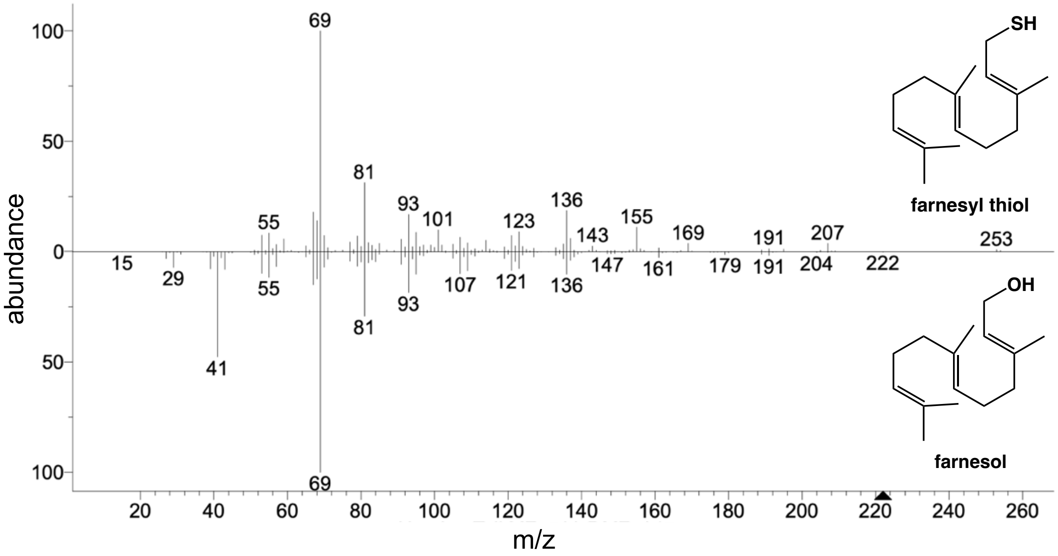


**Figure S4B**. Butterfly plot of mass spectra of a farnesol standard (bottom) and the product produced when AsDMS is incubated with FSPP, interpreted as farnesyl thiol (top). The fragmentation patterns are largely similar but not identical. While a peak at m/z = 238 is not observed, it is thought that the peak is too small to be observed. A small, spurious peak is observed at m/z = 253 corresponding to an impurity; this spurious peak is also observed in the mass spectrum of drimenol in Figure S2. The peak at m/z = 207 results from leaching of the polystyrene stationary phase of the GC column.

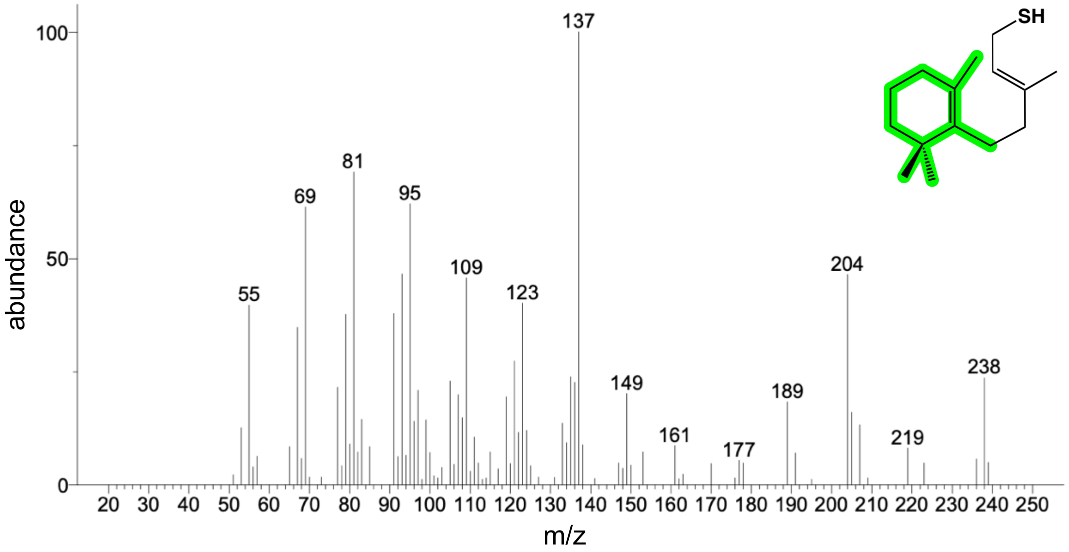


**Figure S4C**. Mass spectrum of unidentified sesquiterpene generated from FSPP along with a proposed structure. Based on the prominent peak at m/z = 238, this compound is a sesquiterpene thiol. Additionally, the robust peak at m/z = 137 could correspond to the monocyclic fragment highlighted in green, which would represent a prematurely quenched intermediate in the cyclization cascade leading to drimenyl thiol.

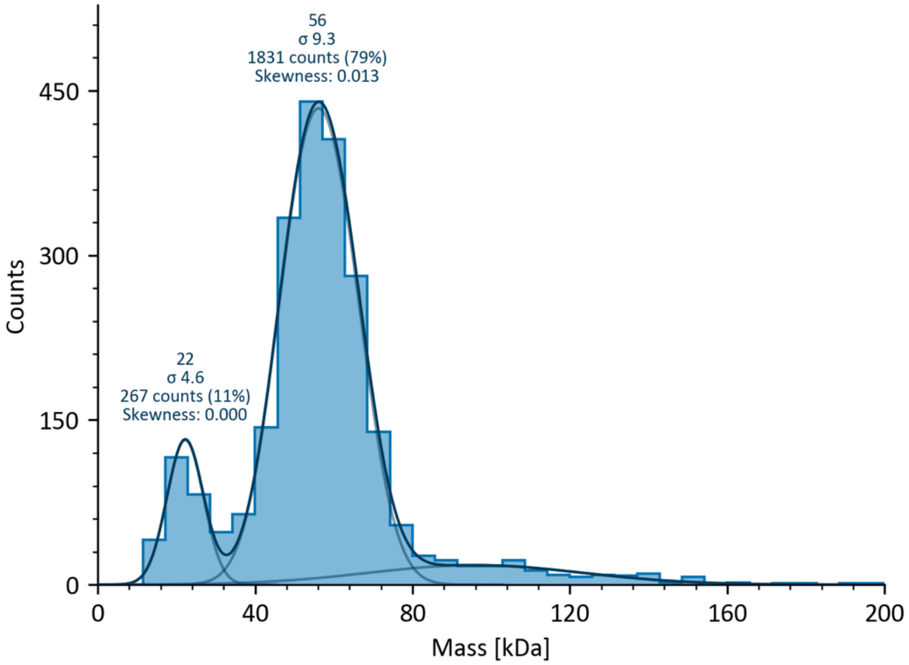


**Figure S5.** Mass photometry measurements indicate that AsDMS is a monomer in solution. The molecular weight of the N-terminal deletion variant prepared for this study (residues 1–526) is 61 kD; the predominant species observed by mass photometry is 56 kD. The low molecular weight species at 22 kD is an artifact.

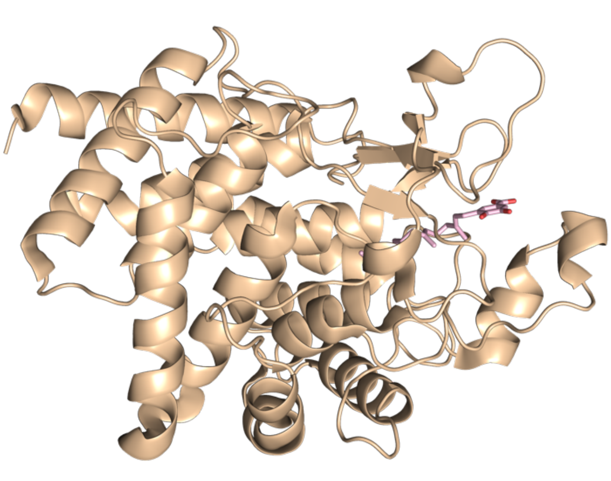

**Figure S6.** Side by side structural comparison of MstE (left) and the β domain of AsDMS (right).
